# Propagation of THz irradiation energy through aqueous layers: Demolition of actin filaments in living cells

**DOI:** 10.1101/846295

**Authors:** Shota Yamazaki, Masahiko Harata, Yuya Ueno, Masaaki Tsubouchi, Keiji Konagaya, Yuichi Ogawa, Goro Isoyama, Chiko Otani, Hiromichi Hoshina

## Abstract

The effect of terahertz (THz) radiation on deep tissues of human body has been considered negligible due to strong absorption by water molecules. However, we observed that the energy of THz pulses transmits a millimeter thick in the aqueous solution, possibly as a shockwave, and demolishes actin filaments. Collapse of actin filament induced by THz irradiation was also observed in the living cells under an aqueous medium. We also confirmed that the viability of the cell was not affected under the exposure of THz pulses. The potential of THz waves as an invasive method to alter protein structure in the living cells is demonstrated.

**One sentence summary:** The energy of the THz wave propagates several millimeters into an aqueous medium and demolishes actin filaments via shockwaves.

## Introduction

Due to the development of terahertz (THz) light sources, industrial and medical applications have been proposed in this decades. Also, toxicity of THz radiation for human health has attracted keen interest among researchers working in this frequency region ^1^. Two projects, the European THz-BRIDGE and the International EMF project in the SCENIHR ^2^, summarize recent studies about THz radiation effects for human body. For example, non-thermal impacts on DNA stability ^3–5^ was induced by THz wave, which could cause chromosomal aberrations in human lymphocytes ^6^. The transcriptional activation of wound-responsive genes in mouse skin ^7^ and DNA damage in an artificial human 3D skin tissue model ^8^ have also been demonstrated. Most of those studies focus on epithelial and corneal cell lines, because THz photons are totally absorbed at the surface of the tissues due to the intense absorbance of liquid water in this frequency region.

However, if the THz radiation is converted to the other type of energy flow which can propagate into water, irradiation of THz wave may cause damage inside the tissues. In fact, the THz photon energy is once absorbed on the body surface, and converted to the thermal and mechanical energies. We recently observed that THz pulses generate shockwaves at the surface of liquid water ^9^. The generated shockwaves propagate several millimeters in depth. Similar phenomena may occur at the human body. The THz-induced shockwaves may induce mechanical stress to the biomolecules and change their morphology. Such indirect effects of the THz irradiation have not been investigated.

To reveal the effect of THz-induced shockwaves to the biological molecules, we focused on morphology of the actin proteins. Actin has two functional forms, monomeric globular (G)-actin and polymerized filamentous (F)-actin. The actin filament forms the elaborate cytoskeleton network, which plays crucial roles in cell shape, motility, and division ^10^. One advantage of using actin is that we can easily obtain enough purified G-actin from tissues ^11^ to reconstruct polymerization reactions *in vitro*. Actin filaments can be directly observed by fluorescence microscopy by staining with silicon-rhodamine (SiR)-actin ^12^. Because actin has pivotal roles in normal and pathological cell function, including transcriptional regulation, DNA repair, cancer cell metastasis, and gene reprogramming ^13–16^, various chemical compounds and regulatory proteins have been analyzed for research and therapeutic purposes ^17^. In this study, we investigated the effect of THz-induced shockwaves on actin filaments in the view of the THz wave induces shockwave propagation in aqueous medium. The morphological changes of actin filaments were employed as evidence of the shockwave operation. The experiments were performed by the actin aqueous solution and living cell with several THz wave energy densities and shockwave propagation lengths. We also confirmed that the viability of the cell was not affected under the exposure of THz pulses. The potential of THz waves as an invasive method to alter protein structure in the living cells is demonstrated.

## Results and Discussion

### Polymerization of actin in aqueous solution

To mimic the effect of THz irradiation on tissue proteins, a 1 mm-thick aqueous solution of actin protein was subjected to THz pulsed irradiation (Fig. 1A). The actin polymerization reaction was induced by adding F-actin buffer to the G-actin solution. During actin polymerization, 80 μJ/cm^2^/micro-pulse THz pulses were applied from the bottom of the dish. Another actin solution was placed next to the sample without THz irradiation as a control. After 30-min of THz irradiation, a portion of the actin solution was collected for the observation with a fluorescence microscope. The irradiated sample (Fig. 1B, right) clearly lost brightness as compared to the control (Fig. 1B, left), and the number of actin filaments with detectable length decreased. The number of actin filaments was decreased about 50% by the THz pulses (Fig. 1C). Figure 1D shows examples of the magnified images of actin filaments. Actin filaments formed straight structures, and morphological deformation such as branching or molecular aggregation was not observed.

**Figure 1.**
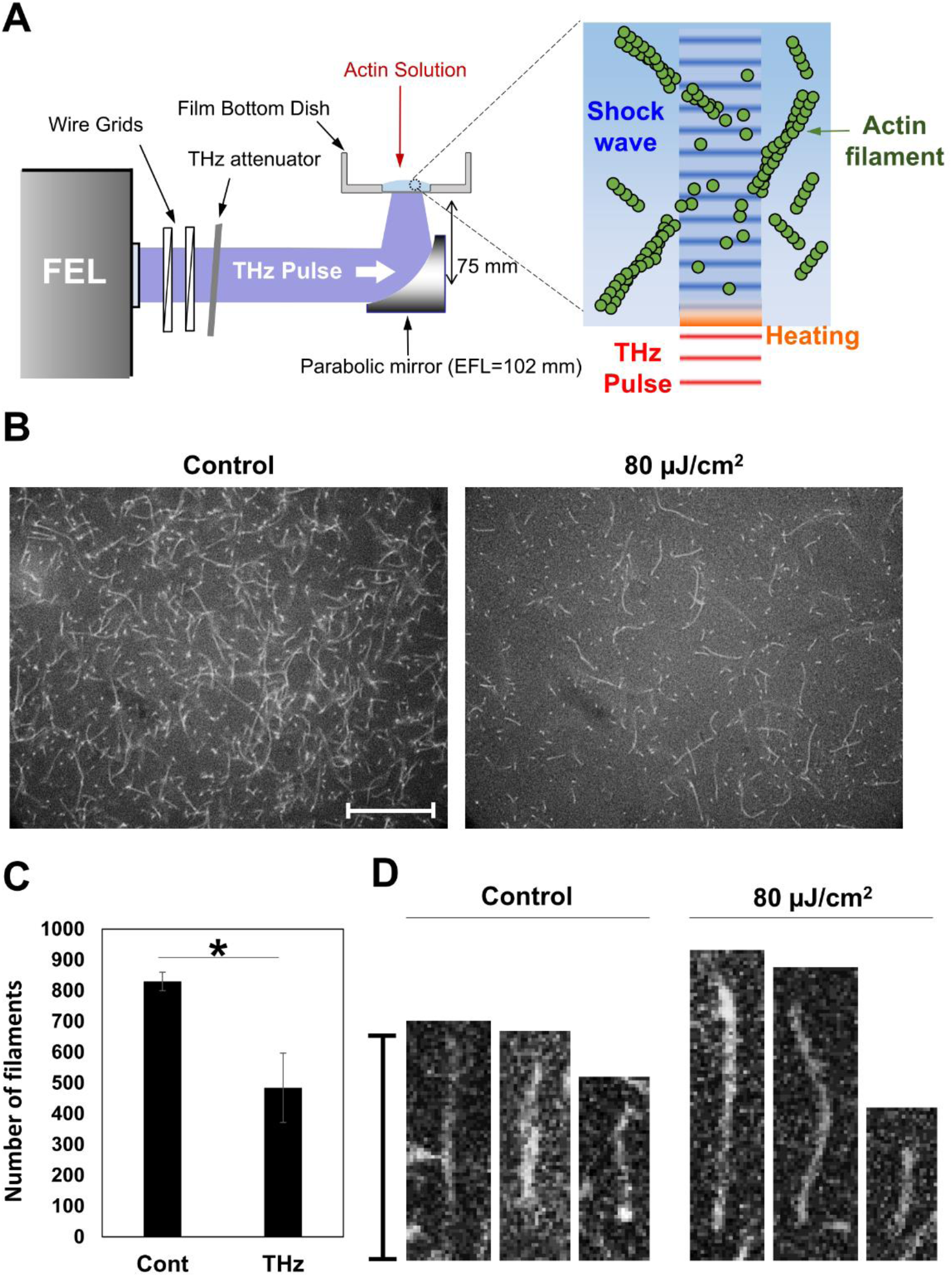
Actin polymerization affected by THz irradiation in an aqueous solution. (A) Schematic diagram of the experimental set-up. Actin solution was placed in a film bottom dish. THz waves (4 THz) were applied to the bottom of the dish. After THz irradiation, a portion of the actin solution was collected for fluorescence microscope observation. (B) Fluorescence microscopy images of actin filaments with (control, left) or without THz irradiation (80 µJ/cm^2^, right). Actin solution (2.4 µM) was fixed and stained with SiR-actin after polymerizing for 30 min. (C) The number of actin filaments counted from fluorescence images. Data shown are the mean and the standard deviation of three independent experiments. More than 300 actin filaments were counted in each of the experiments. Asterisk indicate a statistically significant difference (*P< 0.05). (D) Magnified images of single actin filaments. The bar shows a scale of 20 µm.

When the denaturation of F-actin occurs, an aggregated structure must be observed in the fluorescence image, but aggregation was not observed in fig. 1B. Note that the irradiation of THz pulses to the peptide film was demonstrated using same THz-FEL setup^18^, and denaturation of the sample was observed under the irradiation fluence more than 2 mJ/cm^2^. In contrast, the irradiation fluence in this paper is one order lower than in the previous experiments, and most of the irradiated energy is absorbed by the water molecules in the actin solution. Therefore, we concluded that denaturation of F-actin was not induced in this experiment.

The actin polymerization process consists of three phases: nucleation, elongation, and equilibrium ^19^. In the initial step, four actin monomers combine to form a nucleus. Then, elongation progresses until F-actin reaches equilibrium, where elongation and collapse are balanced. At room temperature, 30 min is sufficient to reach equilibrium. Therefore, decreased actin filaments would be due to a reduced elongation reaction or collapse of actin filaments.

Since the reaction rate of the actin polymerization is temperature-dependent, reduced actin filaments could be explained by increased temperature of water due to absorption of THz waves (Fig. S1A). Using an adiabatic model, temperature was increased at the solution surface by about 0.035 °C with a single micro-pulse (Fig. S1B). The solution surface was then cooled by thermal diffusion in the interval between pulses. Thus, the average temperature gradually increases during irradiation of THz pulses until reaching equilibrium. Sample temperature measured at the end of irradiation was 1.4 °C higher than the control. However, actin polymerization does not show remarkable changes until 50 °C ^20,21^. Increasing the temperature by a few °C does not explain the inhibition of polymerization reaction. Another mechanism must be considered. The next possible explanation for reduced actin polymerization is the direct interaction between THz photons and actin molecules. Since THz photon energy is similar to hydrogen bonding vibration energy, macromolecular structure might be altered by THz excitation. However, the high absorption coefficient of water limits THz wave penetration depth to about 10 μm, which is only 1% of the sample volume. Even with extensive diffusion of actin molecules, the direct interaction of the THz wave with the surface of the reaction solution does not explain the significant difference in the number of actin filaments.

### The effects of THz irradiation on cellular actin filament

When high power THz pulses are focused on a material, various nonlinear photophysical and photochemical processes are induced. We recently observed that THz pulses effectively generate shockwaves, which propagates a few millimeters into the aqueous medium ^9^. As discussed in previously, the temperature change due to the THz irradiation is negligibly small in our experimental conditions. Therefore, shockwave propagation is the most likely reason for THz-induced decrease in actin filaments.

To demonstrate the THz pulse radiation effect deeper into the aqueous layer, we observed morphological changes in actin fibrils from living cells. The cells were sunk in the culture medium to avoid direct interaction with the THz wave and rapid temperature change. HeLa cells were grown on a glass plate and placed 800 µm away from the bottom of the dish to simulate similar depth from a tissue surface. The culture medium was kept at 37 °C during the experiment and THz pulses were applied from the bottom of the dish (Fig. S2). As a control, the same experiment was performed by cutting off the THz beam. The temperature at the center of the glass plate was measured at the end of each experiment. The temperature difference between control and irradiated samples was less than 0.5 °C. Thermal shock in cells does not occur until 42 °C. Therefore, cellular heating by the THz irradiation is negligible in this experiment.

In HeLa cells, actin filaments form several assemblies categorized as cell cortex fibers, stress fibers, and cytoplasmic actin filaments, which are present in different cellular regions. To observe morphological changes, endogenous actin filaments were observed with a fluorescent microscope. HeLa cells were stained with SiR-actin for live-cell imaging before (Fig. 2A, i-iv) and after 30-minute THz irradiation (Fig. 2A, v-viii). The energy of the THz micro-pulse was 0, 80, 160, 250 μJ/cm^2^, respectively. Intact actin filament structure is clearly observed at before THz irradiation in all samples (Fig. 2A, i-iv). This structure was maintained after 30 minutes in control cells (Fig. 2A, v), showing that no significant damage occurs from the UV light of the microscope. After THz irradiation, actin filaments were decreased and actin aggregates (multiprotein complex) were observed in the cell cortex (Fig. 2A, vi-viii, red arrow), which is a layer of thin actin filaments formed at the cell periphery and plays important roles in cell morphology, migration, and invasion.

**Figure 2.**
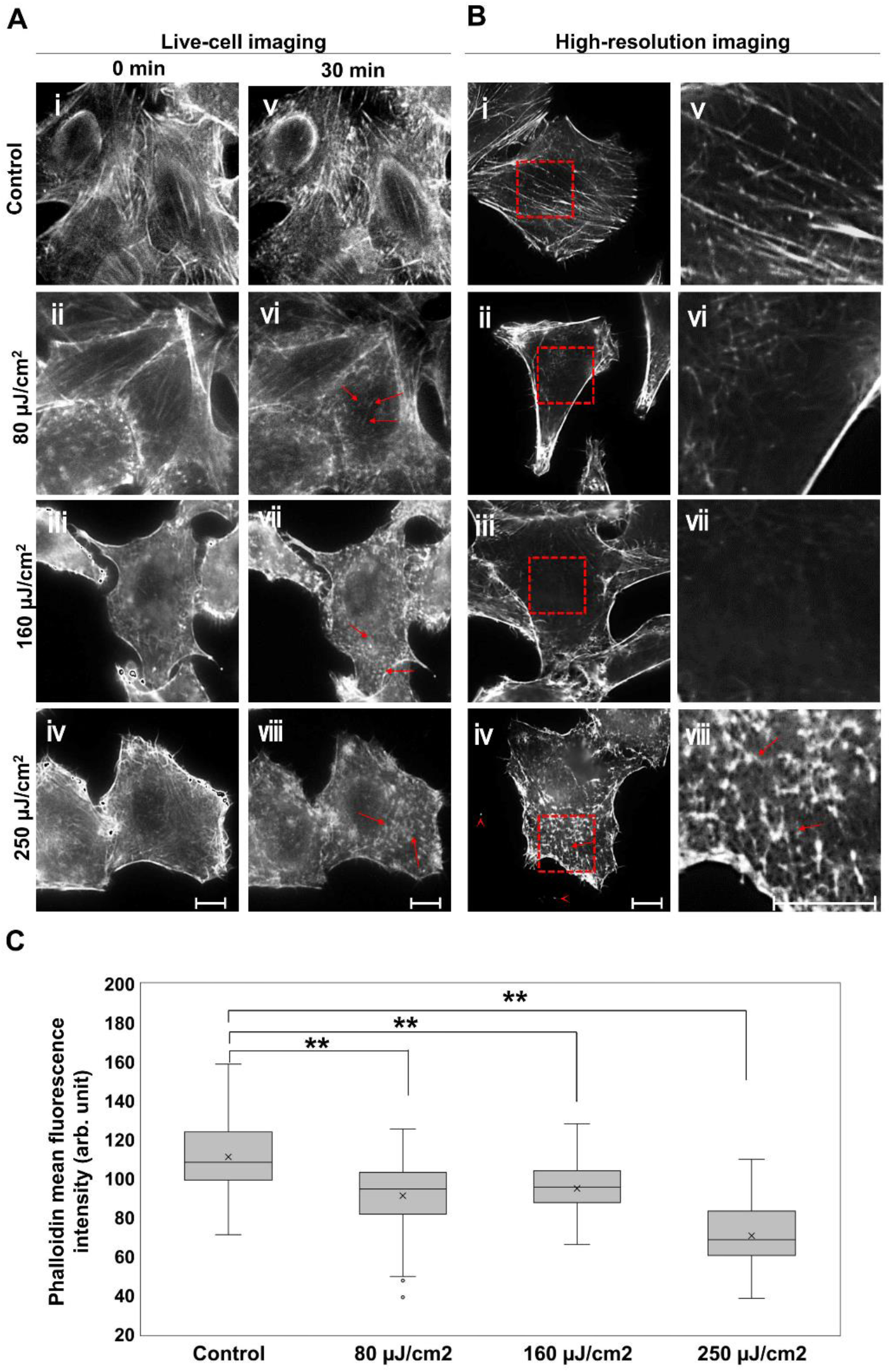
THz wave effects on cellular actin filaments. (A) Live-cell imaging of actin filaments. The energy of the THz irradiation was 0, 80, 160, and 250 µJ/cm^2^. Cells imaged before (A, i-iv) and after 30-minute THz irradiation (A, v-viii). The red arrows indicate aggregated actin filaments. (B) Immunofluorescence images of the cell cortex stained with AlexaFluor 594-phalloidin. Images were observed using spinning-disk confocal microscopy. The right panels (v-viii) show the magnified images of the red squares in the left panels (i-iv), respectively. Red arrows indicate aggregated actin filaments. Arrowheads indicate filipodia fragments. Actin filaments were stained with SiR-actin and observed by fluorescence microscopy. The bar shows a scale of 10 µm. (C) Mean fluorescence intensity of phalloidin was analyzed using immunofluorescence images of the cell cortex. N >51 cells/group. Asterisks indicate a statistically significant difference (\*\**P* < 0.01).

To observe detailed actin filament structure within the cell cortex, cells were immunostained with AlexaFluor 594 Phalloidin and observed by spinning-disk confocal microscopy after THz irradiation for 30 minutes (Fig. 2B, i-iv and v-viii (zoomed images)) (Fig. S3). The samples after 80 and 160 μJ/cm^2^ THz irradiation show a dark area in the cell cortex (Fig. 2B, vi and vii), indicating disassembly of the actin filaments. Actin aggregation is present near the cell periphery (Fig. 2B, iv and viii, red allow) after 250 μJ/cm^2^ irradiation. We also observed severed filopodia in these cells (Fig. 2B, iv, red allow head). Artificial collapse of actin filaments with chemical reagents causes aggregation of actin filaments ^22^, which is similar to the phenomenon shown in Fig. 2B, iv. Figure 2C show the mean fluorescence intensity of Phalloidin in the cell. The mean values were analyzed using immunofluorescence images of the high-resolution imaging. After THz irradiation of 80, 160 and 250 μJ/cm^2^, fluorescence of Phalloidin was reduced in a power-dependent manner. These results demonstrate that actin filaments are disrupted with high THz irradiation power.

### The energy transport of THz wave in water

To examine the energy transport in water induced by THz irradiation, we observed morphological changes in actin fibrils at various distances from the irradiating point (Fig. 3). Cells were positioned 800, 1800, and 2800 μm away from the interface. Similar to Fig. 2, cells were imaged before and after THz irradiation with 250 μJ/cm^2^ micro-pulse energy (Fig. 3A). Figures 3B and S4 show high-resolution images of the cell cortex after irradiation for 30 minutes. Note that the images at 800 μm are taken from Fig. 2 for comparison. The live-cell images at 1800 μm show that the actin filaments disappeared and actin aggregated (Fig. 3A, vii), similar to images at 800 μm (Fig. 3A, vi), which is also present in the high-resolution images (Fig. 3B, vii, red arrow). However, aggregate size is smaller than at 800 μm. At 2800 μm, actin aggregation is not clearly observed (Fig. 3A, viii) and large peripheral actin filaments remain after irradiation (Fig. 3B, viii). Fig. 3C show the mean fluorescence intensity of Phalloidin in the cell, which were positioned at 800, 1800, and 2800 μm. After THz irradiation of 250 μJ/cm^2^ for 30 min, fluorescence of Phalloidin was significantly reduced at 800 and 1800 μm.

**Figure 3.**
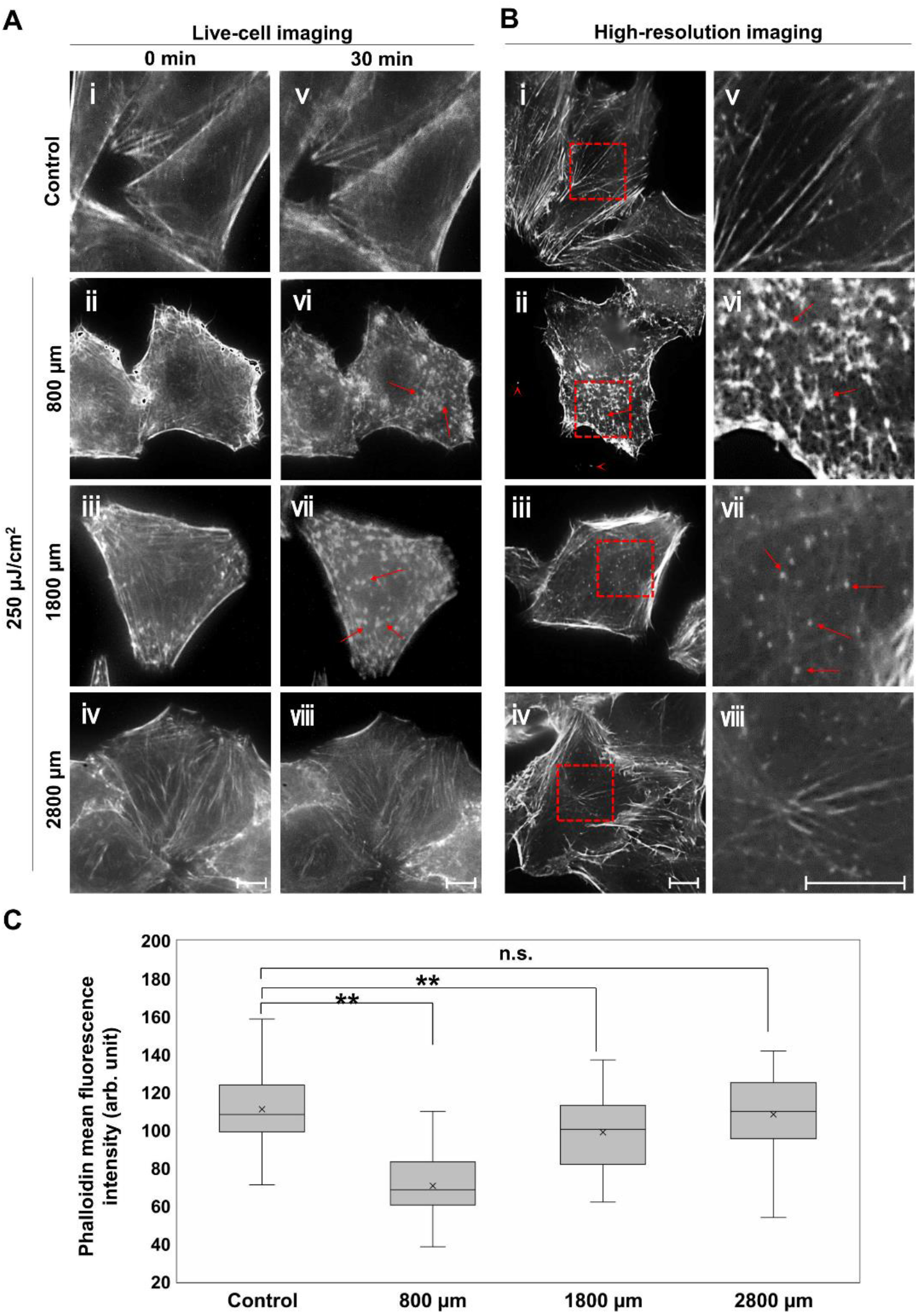
Propagation of THz energy through aqueous culture medium. (A) Live-cell imaging of actin filaments 800, 1800, and 2800 μm away from the bottom of dish. Control cells were placed 800 μm above the dish bottom. Cells imaged before (A, i-iv) and after (A, v-viii) THz radiation with the energy of 250 μJ/cm^2^ for 30 minutes. (B) Immunofluorescence images of the cell cortex stained with AlexaFluor 594-phalloidin. Images were observed using spinning-disk confocal microscopy. The right panels (v-viii) show the magnified images of the red squares in the left panels (i-iv), respectively. Red arrows indicate aggregated actin filaments. Note that the images at 800 μm and control are taken from Fig. 2 for comparison. The bar shows a scale of 10 µm. (C) Mean fluorescence intensity of phalloidin was analyzed using immunofluorescence images of the cell cortex. N >51 cells/group. Asterisks indicate a statistically significant difference (\*\**P* < 0.01).

These results demonstrate that the energy of the THz wave propagates more than 1000 μm into an aqueous solution, which means that the irradiated photon energy is converted to pressure energy and demolishes cellular actin filaments in culture. This result corroborates previous reports which observed that shockwaves generated by piezoceramic vibrator induces actin filament fragmentation ^23,24^. The shockwaves with 16 MPa peak pressure are focused on culture cell and destroys the cell cortex and filopodia, similar to our results (Figs. 2 and 3). Thus, shockwaves generated by THz irradiation propagate deeply into biological tissues, which may change cellular protein morphology.

Recently-developed technologies enable bright, compact, and cheap THz sources. As a result, many applications have been proposed, such as THz body scanners for security applications, next-generation wireless communication, and diagnostic cancer imaging ^25–27^. In the near future, THz waves will become a popular tool used in daily life. However, focused THz radiation might exceed the 80 μJ/cm^2^ energy threshold, changing actin structure. Therefore, the biological effects of THz radiation via shockwaves must be considered when defining safety standards. The effects of THz radiation on surface of the skin have been studied and well summarized ^2^. However, the THz shockwaves penetrated up to a few millimeters into samples. Therefore, information about how deep tissues may be affected is also required when considering the health effects of THz radiation.

### Cytotoxicity of THz irradiation

Shockwaves are used for medical and biological applications, such as surgery, gene therapy, and drug delivery. Near-IR(NIR) and mid-IR (MIR) pulsed lasers have been used for the generation of shockwave. The shockwave generation by NIR and MIR lasers requires the introduction of an absorber or production of the plasma in water or tissues, which are invasive processes with serious risks of damaging biological tissues.^23,24,28,29^. In comparison to the conventional methods via absorbers or plasma generation using IR pulsed lasers, THz light can effectively generate the shockwave in water due to strong absorption via a stretching vibration mode of the hydrogen bonding network. The linear absorption of water is a mild and non-destructive process. Therefore, the proposed method using THz light can be applied to the biological tissue and fragile instruments.

To confirm the “softness” of shockwave generation via THz irradiation, cell death was measured by a cytotoxicity assay using 4’,6-diamidino-2-phenylindole (DAPI). When cell death or loss of membrane integrity occurs, DAPI is taken into the cytoplasm through the damaged membrane and generates fluorescence after DNA binding. Figure 4A shows cells with or without THz irradiation. After THz irradiation for 30 min, cell size was significantly decreased (Fig. S5). Figure 4B shows fluorescence images of DAPI. Figure 4C shows merged image between visible cells and DAPI fluorescence. In contrast to cell shape change, less than 1% of cells show fluorescence after THz irradiation, similar to control experiments. These results indicate that membrane injury or cell death is not induced by THz irradiation, but that manipulation of actin filaments in living cells might be induced by THz waves as shown in Figs. 2 and 3. The demolition of actin filaments by THz irradiation suggests the potential to optically control cellular functions. For example, activation of actin polymerization causes increased cancer cell migration and invasion ^15^. Various compounds affecting actin polymerization have been analyzed for research and therapeutic purposes ^17^. However, it is difficult to derive compounds to cancer cells which localize in deep tissue. We demonstrate that THz irradiation propagates as shockwaves in aqueous solution up to several millimeters in depth. Thus, THz-induced shockwaves could provide precise, chemical-free methods for cancer treatment.

**Figure 4.**
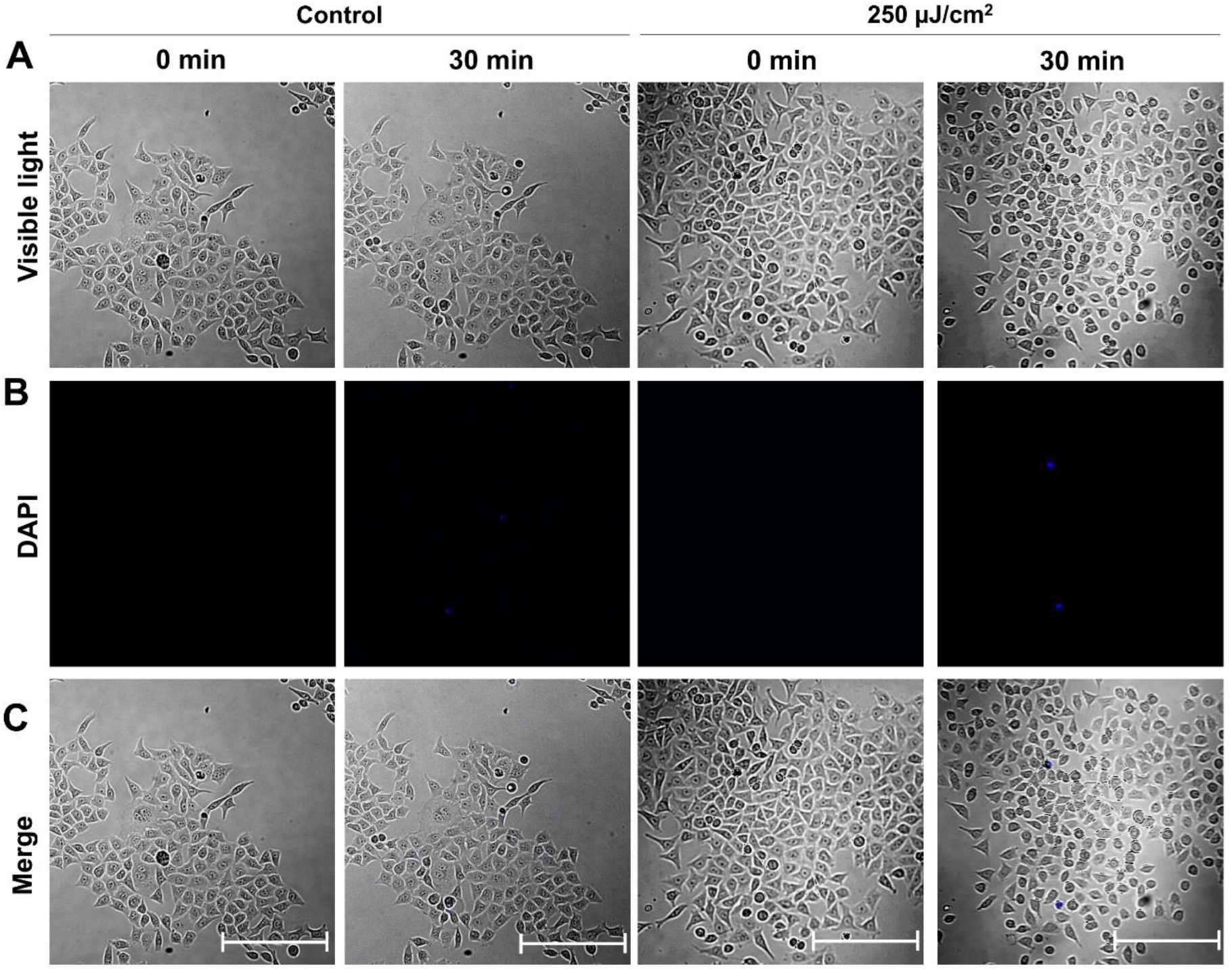
Evidence for no-cytotoxicity by THz irradiation. (A) Visible images of cells. (B) Fluorescence images of DAPI-stained cells (see Material and Methods). (C) Merged image of visible cells and DAPI fluorescence. 0.1 µg/mL DAPI was added to the culture medium at 0 min to detect cell death. The applied THz wave power density was 250 μJ/cm^2^. The cells were placed 800 µm away from the bottom of dish. The bar shows a scale of 100 µm.

## Conclusion

In summary, we demonstrate that THz pulses change actin filament morphology in aqueous solutions via shockwaves. THz irradiation affects not only the surface of the human body, but also a few millimeters into the tissue. This penetration should be considered when developing high-power THz radiation safety standards. These results also indicate that THz irradiation could be applicable for nondestructive manipulation of cellular functions by modulation of actin filaments. For technological development of THz frequency, further work is required to determine the biological kinetics, which is affected by THz irradiation.

## Materials and Methods

### Estimation of THz heating effect with adiabatic model

Here we estimate the temperature rise of water by a single THz pulse by adiabatic model. Figure S1A shows the THz wave irradiated to the water with minute thickness *dx* and surface area *S. I* is the energy of the irradiated THz wave [J].

According to the Lambert Beer’s law, the absorbed THz energy *ΔI* [J] is written as follows,

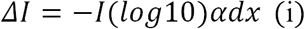

where *α* [cm^-1^] is the absorption coefficient of the liquid water. The increase of the temperature *ΔT* [°C] is

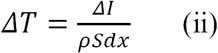

where *ρ* [J/°C cm^3^] is the specific heat of the liquid water. Introducing pulse energy density *P* [J/cm^2^] by the relation *I = P*…*S, ΔT* can be written as follows.

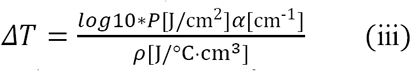

By taking values of *α*=800 cm^-1^, *ρ* = 4.2 J/°C cm^3^, *ΔT* is estimated as 0.035 °C by the irradiation of 80 μJ/cm^2^.

The irradiated pulse is exponentially decayed in the water. Figure S1B shows *ΔT* with a single micro pulse of 80 μJ/cm^2^ irradiated to the surface of the water.

### THz source

A THz free electron laser (FEL) at the Institute of Scientific and Industrial Research (Osaka University) was used as the THz beam source. Details on the THz-FEL were described previously *(20, 21)*. The THz-FEL output was a macro-pulse with 5-Hz repetition, composed of ∼100 micro-pulses with 5-ps duration. The THz wavelength, tuned by the wiggler gap distance, was centered at 4 THz with 1 THz bandwidth. The schematic diagram of the experimental set-up is shown in Fig. 1A. The THz beam was loosely focused by an off-axis parabolic mirror (102-mm focal length), and the sample was offset from the focal point by 25 mm. The beam diameter at the sample was 4 mm. THz power density was regulated with a THz attenuator (CDC Corp: TFA-4) and a pair of wire-grid polarizers. The THz wave energy in front of the sample was estimated by measuring macro-pulse energy with a pyroelectric detector (Coherent Inc.: J-25MB-LE).

### THz irradiation of the actin solution and polymerization reaction

Purified actin protein (Cytoskeleton, Inc.) was used for the *in vitro* actin polymerization reaction. G-actin dissolved into G-buffer (5 mM Tris-HCl pH 8.5, 0.2 mM CaCl_2_) at 0.4 mg/ml and placed on ice for 1 h. Then the solution was centrifuged at 14,000 rpm at 4 °C for 30 min to exclude polymerized actin. The supernatant was used for irradiation experiments. Polymerization was initiated by adding 10× F-buffer (500 mM KCl, 20 mM MgCl_2_, 50 mM guanidine carbonate, and 10 mM ATP). Solution containing 2.4 µM actin was put on an olefin-based film dish (14 mm radius and 1 mm thickness, Matsunami Glass). Polymerization was initiated by adding F-buffer to the actin solution, and started THz irradiation. THz waves were vertically applied from the bottom of the dish (Fig. 1A). THz wave transmittance at the bottom of the dish was 80 % at 4 THz. Actin filament structurs was observed by SiR-actin probe (Cytoskeleton, Inc.) *(12)*. After THz irradiation, sample temperature was measured by a K-type thermocouple (Rixen, TK-6200). Then, 1 µl actin solution was collected from the film dish after stirring and mixed with 9 µl 5.6 µg/ml SiR-actin. Next, 1 µl stained sample was mixed with 3 µl Vectashield mounting medium (Vector Laboratories) and mounted on a slide. The sample was observed with an Olympus IX83 fluorescence microscope. Images were captured with a digital CMOS camera (Hamamatsu, Model C11440-42U30). The number of actin filaments was counted in images from three independent experiments (140 µm×140 µm) using Image J software. Statistical significances were calculated by F- and T-test for the transcriptional assay.

### THz irradiation of HeLa cells

HeLa cells were seeded on 0.15 mm-thick cover glass and cultured in Dulbecco’s modified Eagle’s medium (Gibco) supplemented with 10% fetal bovine serum and antibiotics (penicillin and streptomycin) at 37 °C in 5% CO_2_ humidified atmosphere. Actin filaments were stained with SiR-actin by adding probes from a 1 mM DMSO stock solution to the growth medium (final concentration: 3 µM) and incubating for 1 h at 37 °C in 5% CO_2_ humidified atmosphere. Note that actin organization is not changed by staining *(23)*.

Figure S2 shows the schematic diagram of the experimental set-up for THz irradiation. The cells attached to cover glass were placed in film-bottom dishes filled with culture media. To maintain distance from the bottom of the dish, the cover glass was suspended upside-down as shown in Fig. S2. The distance from the bottom is adjusted by inserting metal spacers. For cytotoxicity analysis, DAPI (0.1 µg/mL) was added to medium at time point 0 min as shown in Fig. 4. We used a commercially available chemical, -Cellstain-DAPI solution, provided by DOUJINDO. This chemical dose not penetrate the intact membrane of mammalian cell and can be used for analysis of cell viability.

The film bottom dish was set on a heating stage (LINKAM: 10002L) to maintain the culture media temperature at 37 ± 1 °C. The THz beam passed vertically through a 4-mm hole in the heating stage. During THz irradiation, fluorescence microscopy images were obtained with a UV light source (Thorlabs, X-Cite 200DC lamp), dichroic mirror (Thorlabs, DMLP650R), two optical filters (excitation 625/25; emission long pass 675), objective lens (Olympus, LUMFLN60XW; Nikon, N10X-PF), and a sCMOS camera (Thorlabs, CS2100M-USB). The cover glass temperature was monitored by a K-type thermocouple after irradiation.

To observe the cell cortex at high resolution, confocal microscopy images were captured. After the irradiation experiment, cells were fixed with 4% paraformaldehyde/phosphate-buffered saline (PBS) and permeabilized with 0.5 % Triton X-100 in PBS for 10 min. Cells were stained with AlexaFluor 594 Phalloidin (Invitrogen) in PBS for 1 h at room temperature. Slides were mounted in Vectasheld mounting medium (Vector Laboratories). Fluorescent microscope images were captured with a fluorescence microscope equipped with a scanning disk unit (Olympus, IX83, IX3-DSU). Image analysis was performed using Fiji software. To measure mean signal intensity in the membrane compartment, the outline of the cell was selected using the area selection tools. The mean signal intensity of the signal over the area of the cell was recorded. The number of cells is shown as *n*. Statistical significances were calculated by F- and T-test for the transcriptional assay.

### THz light induced shockwave

THz light resonates with the intermolecular vibration in the hydrogen-bonding network. The absorption coefficient of water is 800 cm^-1^ at 4 THz *(22)*, which implies that more than 99.7% of irradiated energy is absorbed within 0.1 mm of the water surface. The strong absorption induces a local temperature and pressure increase rapidly. The latter is followed by the shockwave propagation into the water. The shockwave is the wave with the discontinuous change in pressure, and carries the mechanical energy with the speed faster than that of sound.

To demonstrate the THz light induced shockwave, the THz-FEL light was loosely focused to the distilled water in the quartz cell. A spot size of the THz pulse on the water surface was 0.7 mm in diameter. Figure S6 shows a shadowgraph image of the shockwave propagating in the water sample irradiated by the THz-FEL with a micro-pulse energy density of 4.5 mJ/cm^2^ (a power density of 3.1 GW/cm^2^). This image was observed by an image-intensified CCD of the Princeton PI-MAX3 camera with a time gate of 10 ns. A shadowgraph technique clearly displays an inhomogeneous density and pressure distributions in transparent media *(23)*. We can see a stripe pattern in the image which indicates the shockwave propagation observed at the time interval of 36.9 ns. The shadowgraph image indicates that the THz-light-induced shockwave has a plane wave front. The plane shockwave can be generated from a plane source with loosely focused THz-FEL light. The plane wave nature causes the long-distance propagation, more than 3 mm in depth, which is 100 times longer than the skin depth of water for the THz light.

From the snapshot images of the shockwave, we can estimate the speed of the shockwave to be 1491 m/s, which is the sound velocity in distilled water at 23 °C *(24)*.

## Author contributions

S. Y., M. H., Y. O., C. O., and H. H conceived this study. S.Y., Y. U., M.T., and K. K. conducted the experiments and analyzed data. G.I. constructed and managed the THz-FEL facility in Osaka Univ. S.Y., M. T., and H. H. drafted the original manuscript, and all authors edited and reviewed the manuscript.

## Data and materials availability

The data presented in this manuscript are tabulated in the main paper and in the supplementary materials.

## Supplementary Materials

**Figure S1.**
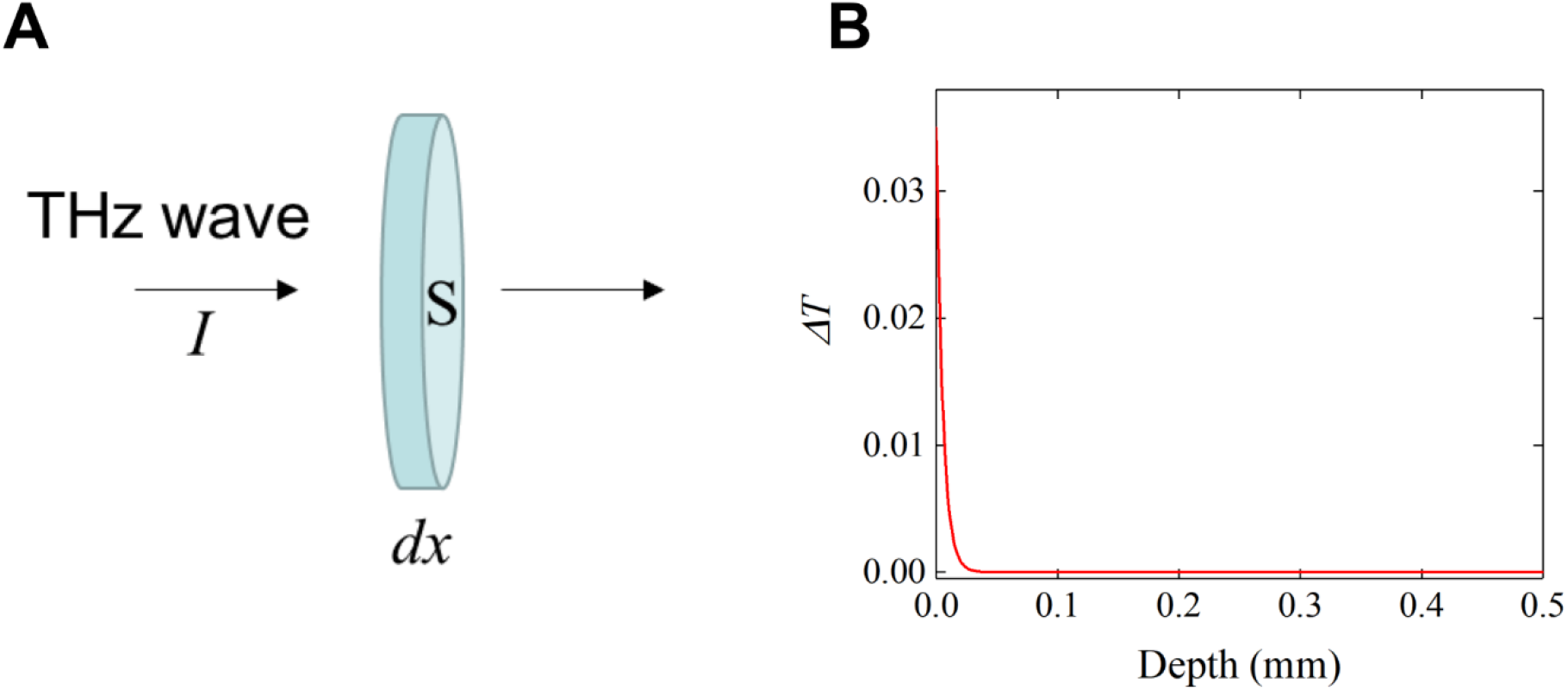
Estimation of THz heating effect with adiabatic model. (A) The model of THz wave irradiated to the water with minute thickness. (B) *ΔT* just after a single micro pulse of 80 μJ/cm^2^ irradiated to the water.

**Figure S2.**
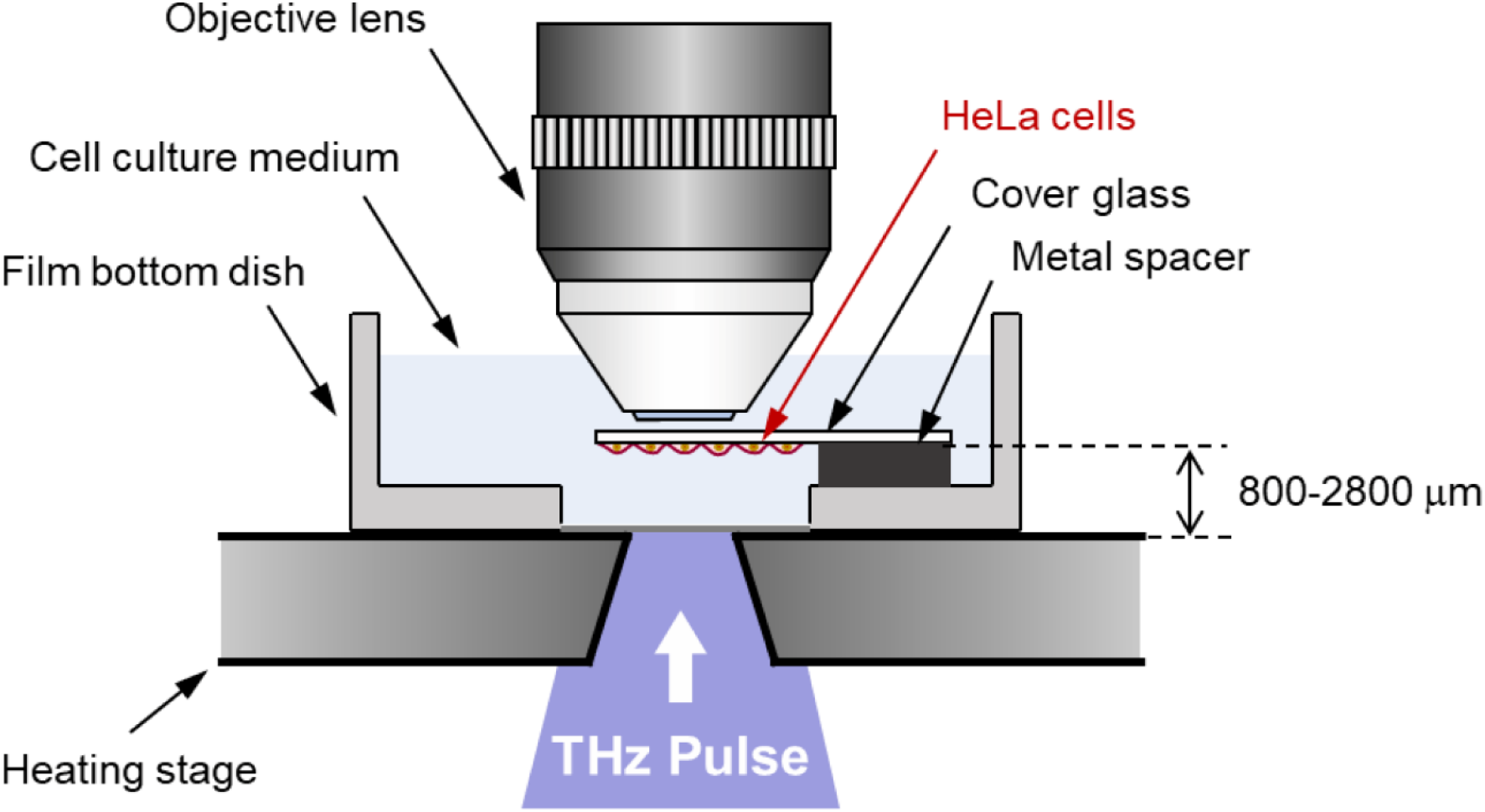
Experimental set-up of cells in culture medium. HeLa cells were seeded on a cover glass and immersed in culture medium. The culture medium was kept at 37 °C during the experiment, and THz pulses were applied from the bottom of the dish.

**Figure S3.**
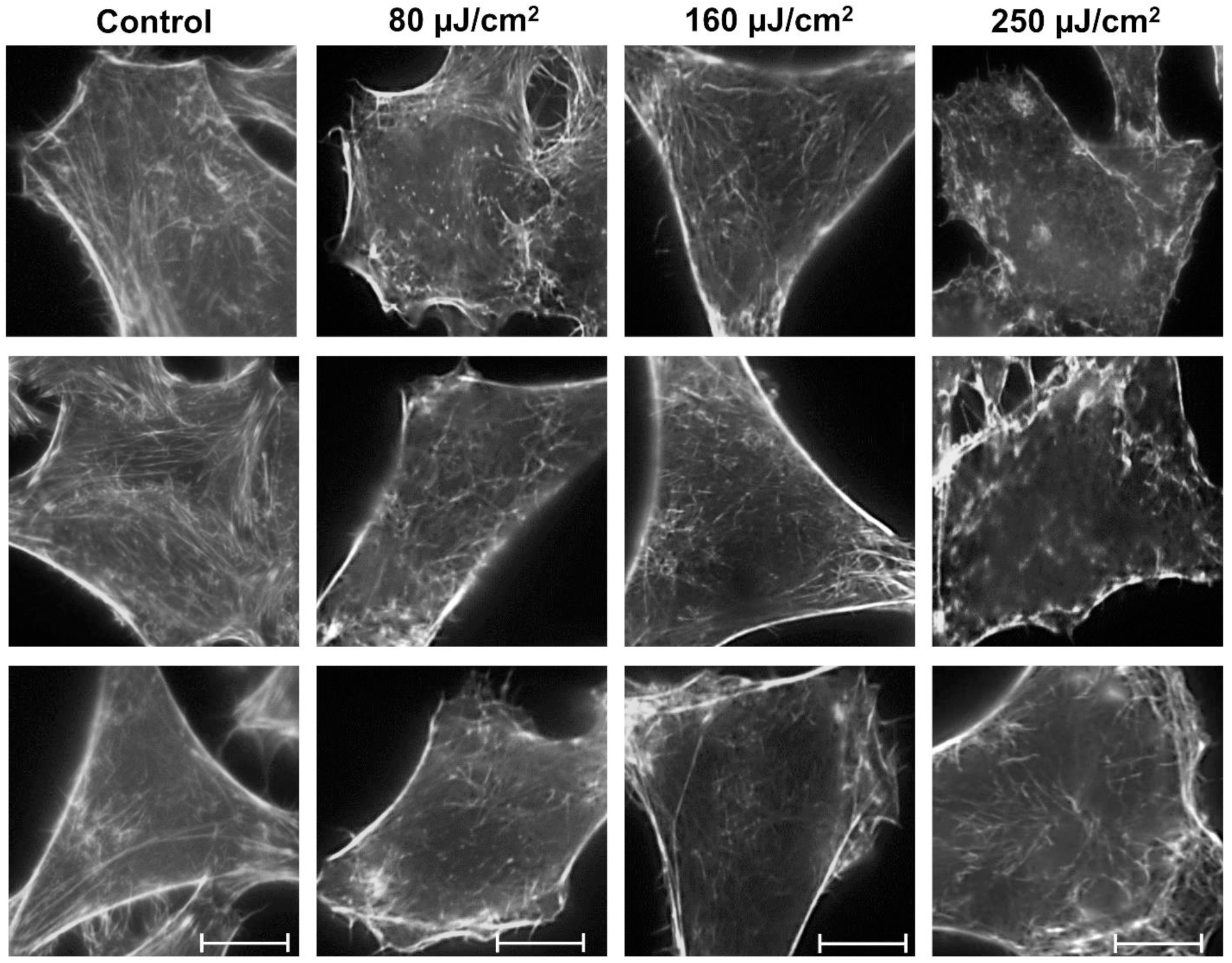
Immunofluorescence images of the cell cortex. After THz irradiation with the indicated energies for 30 min, HeLa cells were fixed and stained with AlexaFluor 594-phalloidin. Images were observed using spinning-disk confocal microscopy. The bar shows a scale of 10 µm.

**Figure S4.**
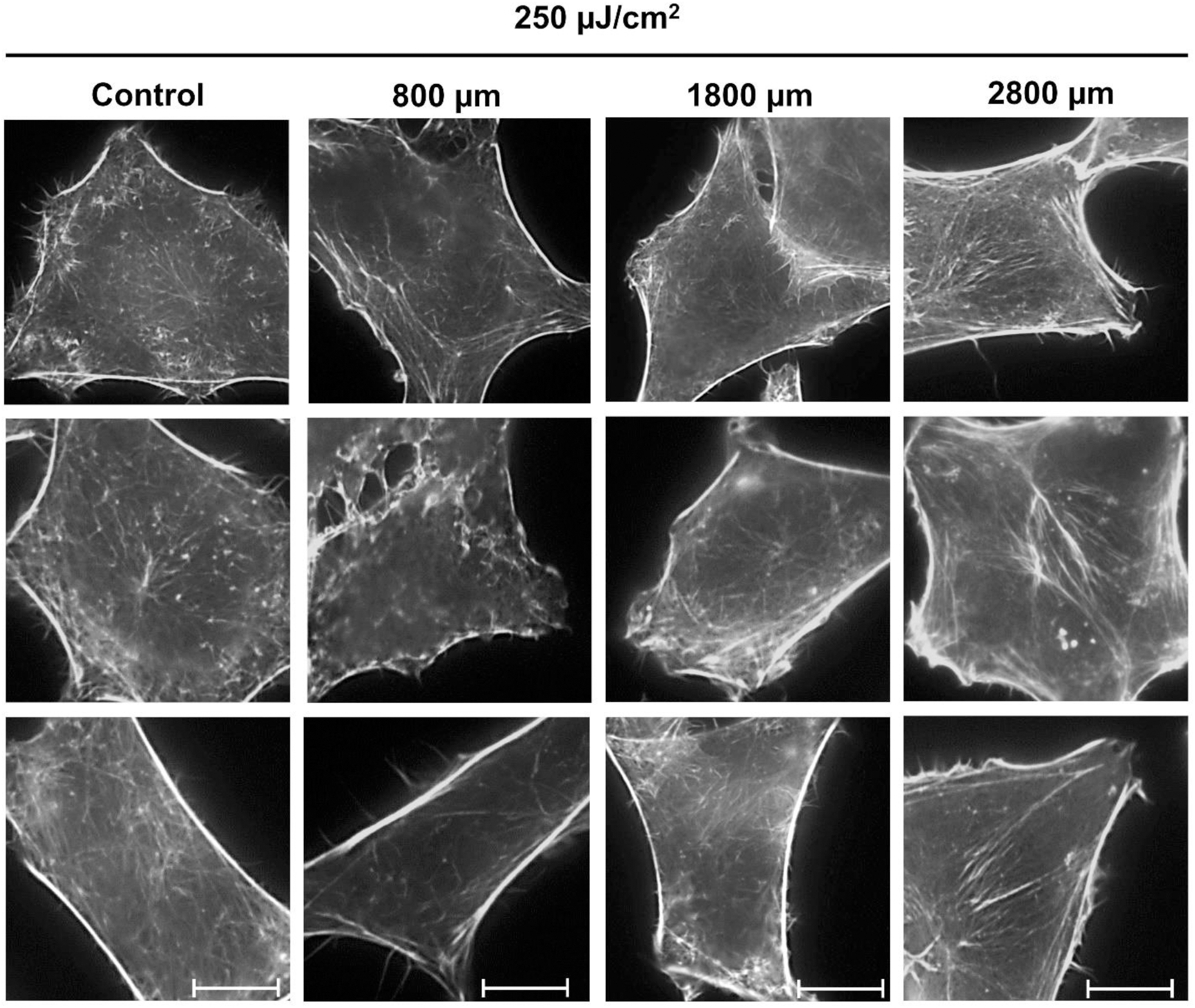
Immunofluorescence images of the cell cortex. After THz irradiation at the indicated distances for 30 min, HeLa cells were fixed and stained with AlexaFluor 594-phalloidin. Images were observed using spinning-disk confocal microscopy. The bar shows a scale of 10 µm.

**Figure S5.**
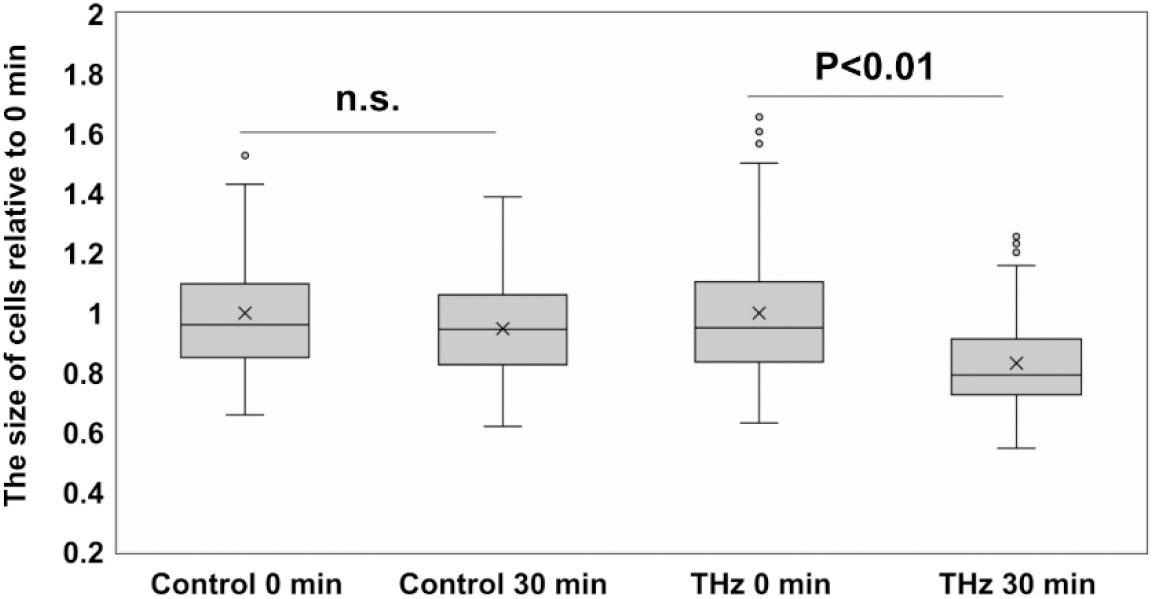
Comparison of cell size before and after THz irradiation. The size of cells was measured in microscopy images using Image J software. The average cell area at 0 min was defined as 1.0. N >80 cells/group.

**Figure S6.**
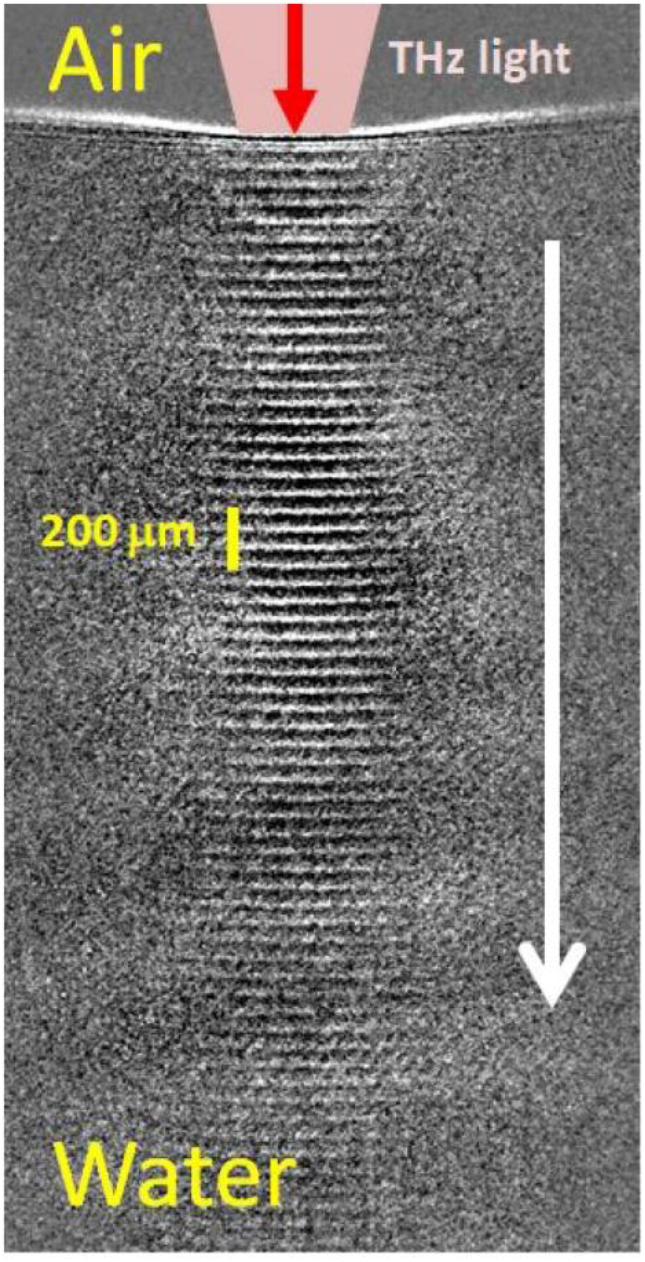
THz-FEL light induced Shockwaves. Snapshot image of a train of shockwaves captured by an image-intensified CCD camera with a time gate of 10 ns. The vertical bar shows a scale of 200 µm.

